# Whole Genome and Exome Sequencing Reference Datasets from A Multi-center and Cross-platform Benchmark Study

**DOI:** 10.1101/2021.02.27.433136

**Authors:** Yongmei Zhao, Li Tai Fang, Tsai-wei Shen, Sulbha Choudhari, Keyur Talsania, Xiongfong Chen, Jyoti Shetty, Yuliya Kriga, Bao Tran, Bin Zhu, Zhong Chen, Wangqiu Chen, Charles Wang, Erich Jaeger, Daoud Meerzaman, Charles Lu, Kenneth Idler, Yuanting Zheng, Leming Shi, Virginie Petitjean, Marc Sultan, Tiffany Hung, Eric Peters, Jiri Drabek, Petr Vojta, Roberta Maestro, Daniela Gasparotto, Sulev Kõks, Ene Reimann, Andreas Scherer, Jessica Nordlund, Ulrika Liljedahl, Jonathan Foox, Christopher Mason, Chunlin Xiao, Wenming Xiao

**Affiliations:** Advanced Biomedical and Computational Sciences, Biomedical Informatics and Data Science Directorate, Frederick National Laboratory for Cancer Research, Frederick, MD, USA; Bioinformatics Research & Early Development, Roche Sequencing Solutions Inc., Belmont, CA, USA; Sequencing Facility, Cancer Research Technology Program, Frederick National Laboratory for Cancer Research, Frederick, MD, USA; Division of Cancer Epidemiology and Genetics, National Cancer Institute, National Institutes of Health, Bethesda, MD, USA; Center for Genomics, School of Medicine, Loma Linda University, Loma Linda, CA, USA; Core Applications Group, Product Development, Illumina Inc, Foster City, CA, USA; Computational Genomics and Bioinformatics Branch, Center for Biomedical Informatics and Information Technology, National Cancer Institute, National Institutes of Health, Bethesda, MD, USA; Computational Genomics, Genomics Research Center (GRC), North Chicago, IL, USA; State Key Laboratory of Genetic Engineering, School of Life Sciences and Shanghai Cancer Center, Fudan University, Shanghai, China; Biomarker Development, Novartis Institutes for Biomedical Research, Switzerland; Companion Diagnostics Development, Oncology Biomarker Development, Genentech, South San Francisco, CA, USA; IMTM, Faculty of Medicine and Dentistry, Palacky University, Olomouc, the Czech Republic; Member of EATRIS ERIC - European Infrastructure for Translational Medicine; Centro di Riferimento Oncologico di Aviano (CRO) IRCCS, National Cancer Institute, Unit of Oncogenetics and Functional Oncogenomics, Italy; Perron Institute for Neurological and Translational Science, Nedlands, Australia; Centre for Molecular Medicine and Innovative Therapeutics, Murdoch University, Murdoch, Australia; Estonian Genome Centre, Institute of Genomics, University of Tartu, Tartu, Estonia; Institute for Molecular Medicine Finland (FIMM), University of Helsinki, Finland; Department of Medical Sciences, Molecular Medicine and Science for Life Laboratory, Uppsala University, Uppsala, Sweden; Department of Physiology and Biophysics, Weill Cornell Medicine, New York, NY, USA, 1305 York Ave., Y13-05, New York, NY, USA; National Center for Biotechnology Information, National Library of Medicine, National Institutes of Health, Bethesda, MD, USA; The Center for Devices and Radiological Health, U.S. Food and Drug Administration, FDA, Silver Spring, MD, USA

## Abstract

With the rapid advancement of sequencing technologies in the past decade, next generation sequencing (NGS) analysis has been widely applied in cancer genomics research. More recently, NGS has been adopted in clinical oncology to advance personalized medicine. Clinical applications of precision oncology require accurate tests that can distinguish tumor-specific mutations from errors or artifacts introduced during NGS processes or data analysis. Therefore, there is an urgent need to develop best practices in cancer mutation detection using NGS and the need for standard reference data sets for systematically benchmarking sequencing platforms, library protocols, bioinformatics pipelines and for measuring accuracy and reproducibility across platforms and methods. Within the SEQC2 consortium context, we established paired tumor-normal reference samples, a human triple-negative breast cancer cell line and a matched normal cell line derived from B lymphocytes. We generated whole-genome (WGS) and whole-exome sequencing (WES) data using 16 NGS library preparation protocols, seven sequencing platforms at six different centers. We systematically interrogated somatic mutations in the paired reference samples to identify factors affecting detection reproducibility and accuracy in cancer genomes. These large cross-platform/site WGS and WES datasets using well-characterized reference samples will represent a powerful resource for benchmarking NGS technologies, bioinformatics pipelines, and for the cancer genomics studies.

## Background & Summary

The NGS technology has become a powerful tool for precision medicine. More researchers and clinicians are utilizing NGS to identify clinically actionable mutations in cancer patients and to establish targeted therapies for patients based on the patient’s genetic makeup or genetic variants of their tumor^1^, there is a critical need to have a full understanding of the many different variables affecting the NGS analysis output. The rapid growing number of sample processing protocols, library preparation methods, sequencing platforms, and bioinformatics pipelines to detect mutations in cancer genome, presents great technical challenges for the accuracy and reproducibility of utilizing NGS for cancer genome mutation detections. To investigate how these experimental and analytical elements may affect mutation detection accuracy, recently we carried out a comprehensive benchmarking study using both whole-genome (WGS) and whole-exome sequencing (WES) data sets generated from two well-characterized reference samples: a human breast cancer cell line (HCC1395) and a B lymphocytes cell line (HCC1395BL) derived from the same donor (NBT-RS47789). We generated WGS and WES data using various NGS library preparation protocols, seven NGS platforms at six centers (NBT-A46164B).

**Figure 1**. shows our overall study design. Briefly, DNA was extracted from fresh cells or cell pellets mimicking the formalin-fixed paraffin-embedded (FFPE) process with fixation time of 1, 2, 6, or 24 hours. A small amount of DNA from fresh cells of HCC1395 and HCC1395BL was pooled at various ratios (3:1, 1:1, 1:4, 1:9 and 1:19) to create mixtures. Both fresh DNA and FFPE DNA were profiled on NGS or microarray platforms following manufacturer recommended protocols. To assess the reproducibility of WGS and WES, six sequencing centers performed a total of 12 replicates (3×3 + 3) on each platform. In addition, 12 WGS libraries constructed using three different library preparation protocols (TruSeq PCR-free, TruSeq-Nano, and Nextera Flex) in four different quantities of DNA inputs (1, 10, 100, and 250 ng) were sequenced on an Illumina HiSeq 4000, and nine WGS libraries constructed using the TruSeq PCR-free protocol were sequenced on an Illumina NovaSeq. Finally, Cytoscan microarray and single-cell sequencing with 10X Genomics platform were performed to uncover the cytogenetics and heterogeneity of two cell lines. Table 1 contains the details of the platform, library protocols and read coverage information.

**Table 1:**
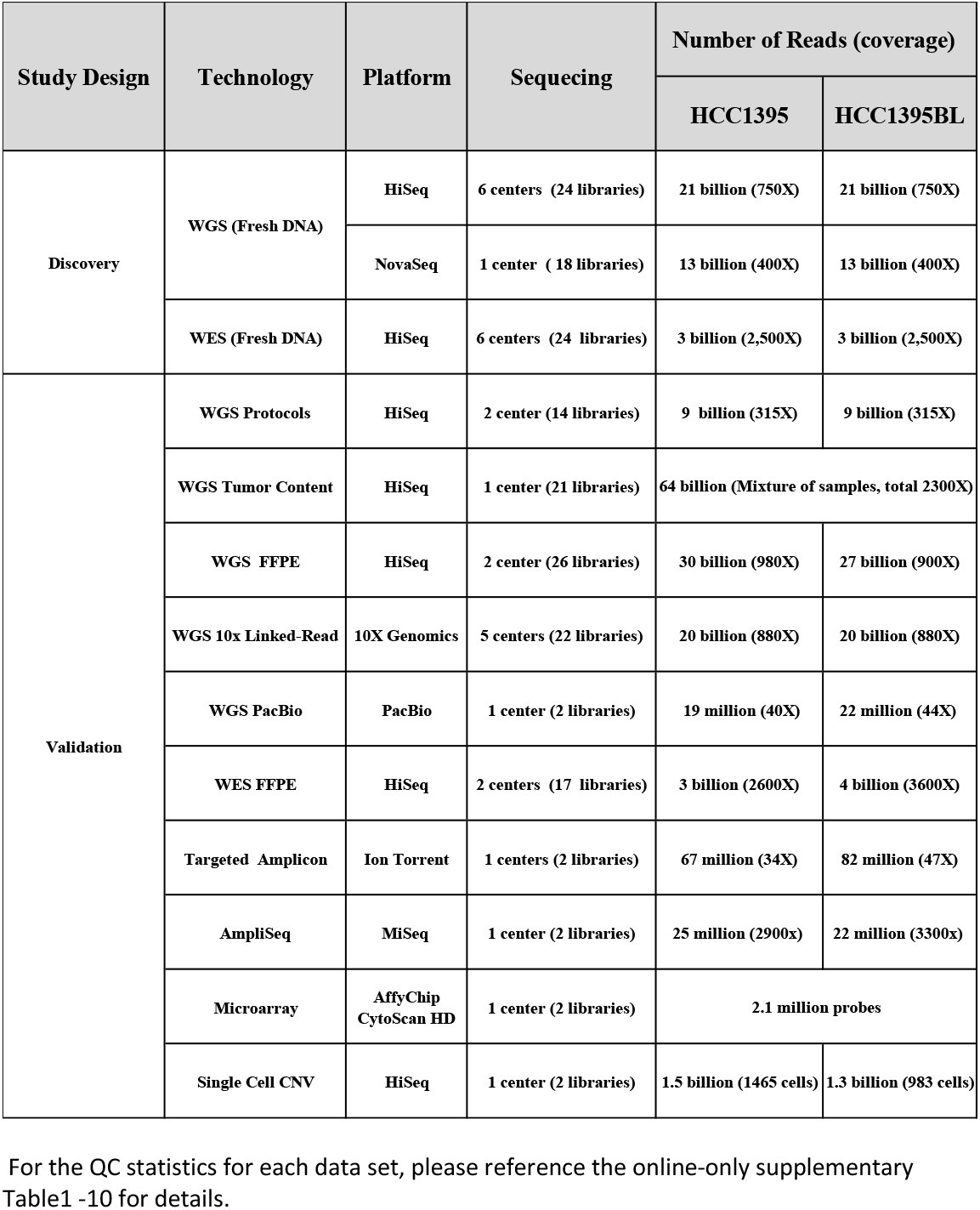
Summary of all experiment data including WGS, WES discovery and validation data sets used in the study.

| Study Design | Technology | Platform | Sequencing | Number of Reads (coverage) |  |
| --- | --- | --- | --- | --- | --- |
|  |  |  |  | HCC1395 | HCC1395BL |
| Discovery | WGS (Fresh DNA) | HiSeq | 6 centers (24 libraries) | 21 billion (750X) | 21 billion (750X) |
|  |  | NovaSeq | 1 center (18 libraries) | 13 billion (400X) | 13 billion (400X) |
|  | WES (Fresh DNA) | HiSeq | 6 centers (24 libraries) | 3 billion (2,500X) | 3 billion (2,500X) |
| Validation | WGS Protocols | HiSeq | 2 center (14 libraries) | 9 billion (315X) | 9 billion (315X) |
|  | WGS Tumor Content | HiSeq | 1 center (21 libraries) | 64 billion (Mixture of samples, total 2300X) |  |
|  | WGS FFPE | HiSeq | 2 center (26 libraries) | 30 billion (980X) | 27 billion (900X) |
|  | WGS 10x Linked-Read | 10X Genomics | 5 centers (22 libraries) | 20 billion (880X) | 20 billion (880X) |
|  | WGS PacBio | PacBio | 1 center (2 libraries) | 19 million (40X) | 22 million (44X) |
|  | WES FFPE | HiSeq | 2 centers (17 libraries) | 3 billion (2600X) | 4 billion (3600X) |
|  | Targeted Amplicon | Ion Torrent | 1 centers (2 libraries) | 67 million (34X) | 82 million (47X) |
|  | AmpliSeq | MiSeq | 1 center (2 libraries) | 25 million (2900x) | 22 million (3300x) |
|  | Microarray | AffyChip<br>CytoScan HD | 1 center (2 libraries) | 2.1 million probes |  |
|  | Single Cell CNV | HiSeq | 1 center (2 libraries) | 1.5 billion (1465 cells) | 1.3 billion (983 cells) |

For the QC statistics for each data set, please reference the online-only supplementary Table1 -10 for details.

**Figure 1.**
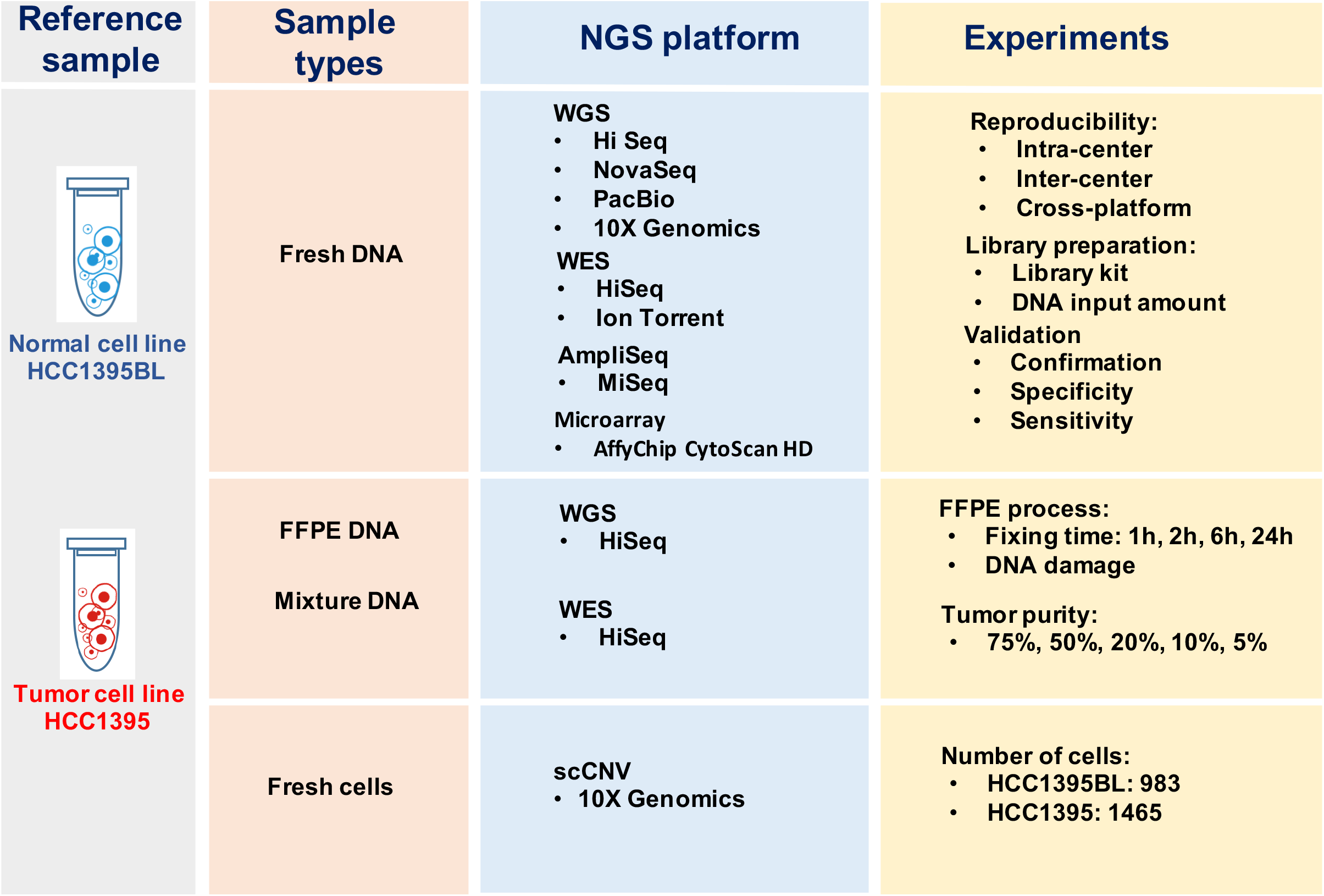
Study design for the experiment. DNA was extracted from either fresh cells or FFPE processed cells. Both fresh DNA and FFPE DNA were profiled on WGS and WES platforms for intra-center, inter-center and cross-platform reproducibility benchmarking. For fresh DNA, six centers performed WGS and WES in parallel following manufacture recommended protocols with limited deviation. Three library preparation protocols (TruSeq-Nano, Nextera Flex, and TruSeq PCR-free,) were used with four different quantities of DNA inputs (1, 10, 100, and 250 ng). DNA from HCC1395 and HCC1395BL was pooled at various ratios to create mixtures of 75%, 50%, 20%, 10%, and 5%. For FFPE samples, each fixation time point (1hm 2h, 6h, 24h) had six blocks that were sequenced at two different centers. All libraries from these experiments were sequenced on the HiSeq series. In addition, nine libraries using the TruSeq PCR-free preparation were run on a NovaSeq for WGS analysis.

We first established reference call sets with evidence from 21 replicates of Illumina WGS runs with coverage ranging from 50X to 100X (1150X in total). We split mutation call confidence levels into four categories: HighConf, MedConf, LowConf, and Unclassified (NBT-RS47789). By combining all WGS runs, we were able to further confirm and improve our call set with tumor-normal pairs of 1500X data sets and identified mutations with VAF as low as 1.5%. A subset of reference mutation calls was validated by targeted exome sequencing (WES at 2,500X coverage) using HiSeq, and deep sequencing from AmpliSeq (at 2,000X coverage) using Miseq, and Ion Torrent (at 34X coverage), and long-read WGS by PacBio Sequel (at 40X coverage). In addition, we inferred subclones and heterogeneity of HCC1395 with bulk DNA sequencing. The results were confirmed by single-cell DNA sequencing analysis (NBT-RS47789B).

With defined reference call sets, we then systematically interrogated somatic mutations to identify factors affecting detection reproducibility and accuracy. By examining the interactions and effects of NGS platform, library preparation protocol, tumor content, read coverage, and bioinformatics process concomitantly, we observed that each component of the sequencing and analysis process can affect the final outcome. Overall WES and WGS results have high concordance and correlation. WES had a better coverage/cost ratio than WGS. However, sequencing coverage of the WES target regions was not even. In addition, WES showed more batch effects/artifacts due to laboratory processing and thus had larger variation between runs, laboratories, and likely between researchers preparing the libraries. As a result, WES had much larger inter-center variation and was less reproducible than WGS. Biological (library) replicates removed some artifacts due to random events (“Non-Repeatable” calls) and offered much better calling precision than did a single test. Analytical repeats (two bioinformatics pipelines) also increased calling precision at the cost of increased false negatives. We found that biological replicates are more important than bioinformatics replicates in cases where high specificity and sensitivity are needed (NBT-RS47789B).

## Methods

Detailed methods were described in our two papers (NBT-A46164B and NBT-RS47789B, in press).

### Cell line culture and DNA extraction

HCC1395; Breast Carcinoma; Human (*Homo sapiens*) cells (expanded from ATCC CRL-2324) were cultured in ATCC-formulated RPMI-1640 Medium, (ATCC 30-2001) supplemented with fetal bovine serum (ATCC 30-2020) to a final concentration of 10%. Cells were maintained at 37 °C with 5% carbon dioxide (CO_2_) and were sub-cultured every 2 to 3 days, per ATCC recommended procedures using 0.25% (w/v) Trypsin-0.53 mM EDTA solution (ATCC 30-2101), until appropriate densities were reached. HCC1395BL; B lymphoblast; Epstein-Barr virus (EBV) transformed; Human (*Homo sapiens*) cells (expanded from ATCC CRL-2325) were cultured in ATCC-formulated Iscove’s Modified Dulbecco’s Medium, (ATCC Catalog No. 30-2005) supplemented with fetal bovine serum (ATCC 30-2020) to a final concentration of 20%. Cells were maintained at 37 °C with 5% CO_2_ and were sub-cultured every 2 to 3 days, per ATCC recommended procedures, using centrifugation with subsequent resuspension in fresh medium until appropriate densities were reached. Final cell suspensions were spun down and re-suspended in PBS for nucleic acid extraction.

All cellular genomic material was extracted using a modified Phenol-Chloroform-Iso-Amyl alcohol extraction approach. Essentially, cell pellets were re-suspended in TE, subjected to lysis in a 2% TritonX-100/0.1% SDS/0.1 M NaCl/10mM Tris/1mM EDTA solution and were extracted with a mixture of glass beads and Phenol-Chloroform-Iso-Amyl alcohol. Following multiple rounds of extraction, the aqueous layer was further treated with Chloroform-IAA and finally underwent RNases treatment and DNA precipitation using sodium acetate (3 M, pH 5.2) and ice-cold Ethanol. The final DNA preparation was re-suspended in TE and stored at -80°C until use.

### FFPE processing and DNA extraction

Please see Online methods in manuscript NBT-RA46164 for details.

### Illumina WGS Library Preparation

The TruSeq DNA PCR-Free LT Kit (Illumina, FC-121-3001) was used to prepare samples for whole genome sequencing. WGS libraries were prepared at six sites with the TruSeq DNA PCR-Free LT Kit according to the manufacturers’ protocol. The input DNA amount for WGS library preparation with fresh DNA for TruSeq-PCR-free libraries was 1 ug unless otherwise specified. All sites used the same fragmentation conditions for WGS by using Covaris with targeted size of 350 bp. All replicated WGS were prepared on a different day.

The concentration of the TruSeq DNA PCR-Free libraries for WGS was measured by qPCR with the KAPA Library Quantification Complete Kit (Universal) (Roche, KK4824). The concentration of all the other libraries was measured by fluorometry either on the Qubit 1.0 fluorometer or on the GloMax Luminometer with the Quant-iT dsDNA HS Assay kit (ThermoFisher Scientific, Q32854). The quality of all libraries was assessed by capillary electrophoresis either on the 2100 Bioanalyzer or TapeStation instrument (Agilent) in combination with the High Sensitivity DNA Kit (Agilent, 5067-4626) or the DNA 1000 Kit (Agilent, 5067-1504) or on the 4200 TapeStation instrument (Agilent) with the D1000 assay (Agilent, 5067-5582 and 5067-5583).

For the WGS library preparation from cross-site study, the sequencing was performed at six sequencing sites using three different Illumina platforms including HiSeq 4000 instrument at 2 x 150 bases read length with HiSeq 3000/4000 SBS chemistry (cat# FC-410-1003), and on a NovaSeq instrument at 2 x 150 bases read length using the S2 configuration (cat#PN 20012860), or on a HiSeq X Ten at 2×150bases read length using the X10 SBS chemistry (cat# FC-501-2501). Sequencing was performed following the manufacturer’s instructions.

For the comparison study of WGS library protocol using different input DNA amounts, Illumina TruSeq DNA PCR-free protocol used 250ng input DNA, Illumina TruSeq Nano protocol libraries were prepared with 1ng, 10ng, and 100ng input DNA amounts. Illumina Nextera Flex libraries were prepared with 1ng, 10ng, and 100ng input DNA amounts. These libraries sequenced at two sequencing sites using two different Illumina platforms including HiSeq 4000 instrument (Illumina) at 2 x 150 bases read length with HiSeq 3000/4000 SBS chemistry (Illumina, FC-410-1003) and NovaSeq instrument (Illumina) at 2 x 150 bases read length using the S2 configuration (Illumina, PN 20012860). Sequencing was performed following the manufacturer’s instructions.

For the tumor purity study, 1µg tumor:normal dilutions were made in the following ratios using Resuspension Buffer (Illumina): 1:0, 3:1, 1:1, 1:4, 1:9, 1:19 and 0:1. Each ratio was diluted in triplicate. DNA was sheared using the Covaris S220 to target a 350 bp fragment size (Peak power 140w, Duty Factor 10%, 200 Cycles/Bursts, 55s, Temp 4 °C). NGS library preparation was performed using the Truseq DNA PCR-free protocol (Illumina) following the manufacturer’s recommendations. The sample purity WGS libraries were sequenced on a HiSeq 4000 instrument (Illumina) at 2 x 150 bases read length with HiSeq 3000/4000 SBS chemistry (Illumina, FC-410-1003). Sequencing was performed following the manufacturer’s instructions.

### Whole Exome Library Construction and Sequencing

SureSelect Target Enrichment Reagent kit, PTN (Part No G9605A), SureSelect Human All Exon v6 + UTRs (Part No 5190-8881), Herculase II Fusion DNA Polymerase (Part No 600677) from Agilent Technologies and Ion Xpress Plus Fragment kit (Part No 4471269, Thermo Fischer Scientific Inc) were combined to prepare library according to the manufacturer’s guidelines (User guide: SureSelect Target Enrichment System for Sequencing on Ion Proton, Version C0, December 2016, Agilent Technologies). Prior, during and after library preparation the quality and quantity of genomic DNA (gDNA) and/or libraries were evaluated applying QubitTM fluorometer 2.0 with dsDNA HS Assay Kit (Thermo Fischer Scientific Inc) and Agilent Bioanalyzer 2100 with High Sensitivity DNA Kit (Agilent Technologies).

WES libraries were sequenced at six sequencing sites with two different Illumina platforms, Hiseq4000 instrument (Illumina) at 2×150 bases read length with HiSeq 3000/4000 SBS chemistry (Illumina, FC-410-1003) and Hiseq2500 (Illumina) at 2×100 bases read length with HiSeq2500 chemistry (Illumina, FC-401-4003). Sequencing was performed following the manufacturer’s instructions.

### Whole Genome FFPE Sample Library Preparation and Sequencing

For the FFPE WGS study, NEBNext Ultra II (NEB) libraries were prepared according to the manufacturer’s instructions. However, input adjustments were made according to the dCq obtained for each sample using the TruSeq FFPE DNA Library Prep QC Kit (Illumina) to account for differences in sample amplifiability. A total of 33 ng of amplifiable DNA was used as input for each sample.

FFPE WGS libraries were sequenced on two different sequencing canters on Hiseq4000 instrument (Illumina) at 2×150 bases read length with HiSeq 3000/4000 SBS chemistry (Illumina, FC-410-1003). Sequencing was performed following the manufacturer’s instructions.

### Whole Exome FFPE Sample Library Preparation and Sequencing

For the FFPE study, SureSelect (Agilent) WES libraries were prepared according to the manufacturer’s instructions for 200 ng of DNA input, including reducing the shearing time to four minutes. Additionally, the adaptor-ligated libraries were split in half prior to amplification. One half was amplified for 10 cycles and the other half for 11 cycles to ensure adequate yields for probe hybridization. Both halves were combined after PCR for the subsequent purification step.

FFPE WES libraries were sequenced on at two sequencing sites with different Illumina platforms, Hiseq4000 instrument (Illumina) at 2×150 bases read length with HiSeq 3000/4000 SBS chemistry (Illumina, FC-410-1003) and Hiseq2500 (Illumina) at 2×100 bases read length with HiSeq2500 chemistry (Illumina, FC-401-4003). Sequencing was performed following the manufacturer’s instructions.

### PacBio Library Preparation and Sequencing

15 ug of material was sheared to 40 kbp with Megarupter (Diagenode). Per the Megarupter protocol the samples were diluted to <50 ng/ul. A 1x AMPure XP bead cleanup was performed. Samples were prepared as outlined on the PacBio protocol titled “Preparing >30 kbp SMRTbell Libraries Using Megarupter Shearing and Blue Pippin Size-Selection for PacBio RS II and Sequel Systems.” After library preparation, the library was run overnight for size selection using the Blue Pippin (Sage). The Blue Pippin was set to select a size range of 15-50 kbp. After collection of the desired fraction, a 1x AMPure XP bead cleanup was performed. The samples were loaded on the PacBio Sequel (Pacific Biosciences) following the protocol titled “Protocol for loading the Sequel.” The recipe for loading the instrument was generated by the Pacbio SMRTlink software v5.0.0. Libraries were prepared using Sequel chemistry kits v2.1, SMRTbell template kit 1.0 SPv3, magbead v2 kit for magbead loading, sequencing primer v3, and SMRTbell clean-up columns v2. Libraries were loaded at between 4 pM and 8 pM.Sequencing was performed following the manufacturer’s instructions.

### 10X Genomics Chromium Genome Library Preparation and Sequencing

Sequencing libraries were prepared from 1.25 ng DNA using the Chromium Genome Library preparation v2 kit (10X Genomics, cat #120257/58/61/62) according to the manufacturer’s protocol (#CG00043 Chromium Genome Reagent Kit v2 User Guide). The quality of the libraries was evaluated using the TapeStation D1000 Screen Tape (Agilent). The adapter-ligated fragments were quantified by qPCR using the library quantification kit for Illumina (KK4824, KAPA Biosystems) on a CFX384Touch instrument (BioRad) prior to cluster generation and sequencing. Chromium libraries were sequenced on a HiSeq X Ten or a HiSeq 4000 instrument at 2 x 150 base pair (bp) read length and using sequencing chemistry v2.5 or HiSeq 3000/4000 SBS chemistry (Illumina, cat# FC-410-1003) across five sequencing sites. Sequencing was performed following the manufacturer’s instructions.

### AmpliSeq library construction and sequencing

AmpliSeq libraries were prepared in triplicate and prepared as specified in the Illumina protocol (Document # 1000000036408 v04) following the two oligo pools workflow with 10 ng of input genomic DNA per pool. The number of amplicons per pool was 1517 and 1506 respectively. The libraries were quality-checked using an Agilent Tapestation 4200 with the DNA HS 1000 kit and quantitated using a Qubit 3.0 and DNA high sensitivity assay kit. The libraries were applied to a MiSeq v2.0 flowcell. They were then amplified and sequenced with a MiSeq 300 cycle reagent cartridge with a read length of 2 ×150 bp. The MiSeq run produced 7.3 Gbp (94.5%) at ≥Q30. The total number of reads passing filter was 47,126,128 reads.

### Whole Exome library Ion Platform Sequencing

SureSelect Target Enrichment Reagent kit, PTN (Part No G9605A), SureSelect Human All Exon v6 + UTRs (Part No 5190-8881), Herculase II Fusion DNA Polymerase (Part No 600677) from Agilent Technologies and Ion Xpress Plus Fragment kit (Part No 4471269, Thermo Fisher Scientific Inc) were combined to prepare libraries according to the manufacturer’s guidelines (User guide: SureSelect Target Enrichment System for Sequencing on Ion Proton, Version C0, December 2016, Agilent Technologies). Prior, during, and after library preparation the quality and quantity of genomic DNA (gDNA) and/or libraries were evaluated applying QubitTM fluorometer 2.0 with dsDNA HS Assay Kit (Thermo Fisher Scientific Inc) and Agilent Bioanalyzer 2100 with High Sensitivity DNA Kit (Agilent Technologies).

For sequencing the WES libraries, the Ion S5 XL Sequencing platform with Ion 540-Chef kit (Part No A30011, Thermo Fisher Scientific Inc) and the Ion 540 Chip kit (Part No A27766, Thermo Fisher Scientific Inc) were used. One sample per 540 chip was sequenced, generating up to 60 million reads with average length of 200 bp.

### 10X Genomics Single Cell CNV library construction, sequencing and analysis

HCC1395 and HCC1395 BL were cultured as described above. 500,000 cells of each culture were suspended in 1 mL suspension medium (10% DMSO in cell culture medium). Cells were harvested the next day for single-cell copy number variation (CNV) analysis via the 10X Genomics Chromium Single Cell CNV Solution (Protocol document CG000153) produces Single Cell DNA libraries ready for Illumina sequencing according to manufacturer’s recommendations. Libraries were sequenced on a HiSeq 4000 instrument at 2 x 150 base pair (bp) read length and using sequencing chemistry v2.5 or HiSeq 3000/4000 SBS chemistry (Illumina, cat# FC-410-1003). Demultiplex BCL from sequencing run and Copy Number Variation analysis were performed using 10X Genomics Cell Ranger DNA version 1.1 software. CNV and heterogeneity visualization analysis was performed via 10X Genomics Loupe scDNA browser.

### Affymetrix Cytoscan HD microarray

DNA concentration was measured spectrophotometrically using a Nanodrop (Life technology), and integrity was evaluated with a TapeStation 4200 (Agilent). Two hundred and fifty nanograms of gDNA were used to proceed with the Affymetrix CytoScan Assay kit (Affymetrix). The workflow consisted of restriction enzyme digestion with Nsp I, ligation, PCR, purification, fragmentation, and end labeling. DNA was then hybridized for 16 hr at 50 °C on a CytoScan array (Affymetrix), washed and stained in the Affymetrix Fluidics Station 450 (Affymetrix), and then scanned with the Affymetrix GeneChip Scanner 3000 G7 (Affymetrix). Data were processed with ChAS software (version 3.3). Array-specific annotation (NetAffx annotation release 36, built with human hg38 annotation) was used in the analysis workflow module of ChAS. Karyoview plot and segments data were generated with default parameters.

### Reference genome

The reference genome we used was the decoy version of the GRCh38/hg38 human reference genome (https://gdc.cancer.gov/about-data/data-harmonization-and-generation/gdc-reference-files; GRCh38.d1.dv1.fa), which was utilized by the Genomic Data Commons (GDC). The gene annotation GTF file was downloaded from the 10X website as refdata-cellranger-GRCh38-1.2.0.tar.gz, which corresponds to the GRCh38 genome and Ensmebl v84 transcriptome.

All the following bioinformatics data analyses are based on the above reference genome and gene annotation.

### Preprocessing and Alignment of WGS Illumina Data

For each of the paired-end read files (i.e., FASTQ 1 and 2 files) generated by Illumina sequencers (HiSeq, NovaSeq, X Ten platforms), we first trimmed low-quality bases and adapter sequences using Trimmomatic^2^. The trimmed reads were mapped to the human reference genome GRCh38 (see the read alignment section) using BWA MEM (v0.7.17)^3^ in paired-end mode and bwa-mem was run with the –M flag for downstream Picard^5^ compatibility.

Post alignment QC was performed based both FASTQ on BWA alignment BAM files, the read quality and adapter content were reported by FASTQC^4^ software. The genome mapped percentages and mapped reads duplication rates calculated by BamTools (v2.2.3) and Picard (v1.84)^5^. The genome coverage and exome target region coverages as well as mapped reads insert sizes, and G/C contents were profiled using Qualimap(v2.2)^6^ and custom scripts. Preprocessing QC reports were generated during each step of the process. MultiQC(v1.9)^7^ was run to generate an aggregated report in html format. A standard QC metrics report was generated from a custom script. The preprocessing and alignment QC analysis pipeline is described in **Suppl. Figure 1a**.

### Preprocessing and Alignment of WES Illumina Data

For each of the paired-end read files generated by Illumina sequencers (HiSeq2500, HiSeq4000 platforms), we first trimmed low-quality bases and adapter sequences using Trimmomatic^2^. The trimmed reads were mapped to the human reference genome GRCm38 (see the read alignment section) using BWA MEM (v0.7.17)^3^ in paired-end mode. We calculated on-target rate based on the percentage of mapped reads that were overlap the target capture bait region file (target.bed). The post alignment QC methods are same as WGS Illumina data pre-processing.

### DNA Damage Estimate for WGS, WES and FFPE Samples

The DNA Damage Estimator(v3)^8^ was used to calculate the GIV score based on an imbalance between R1 and R2 variant frequency of the sequencing reads to estimate the level of DNA damage that was introduced in the sample/library preparation processes. GIV score above 1.5 is defined as damaged. At this GIV score, there are 1.5 times more variants on R1 than on R2. Undamaged DNA samples have a GIV score of 1.

### Preprocessing and Alignment of PacBio Data

PacBio raw data were merged bam files using SMRTlink tool v6.0.1. which used minimap2^9^ as default aligner. Duplicate reads were mark and removed from PBSV alignment bases on the reads coming from the same ZMW, the base pair tolerance was set to 100bp to remove the duplicated reads. The preprocessing and alignment QC analysis pipeline for PacBio data is described in **Suppl. Figure 1b**.

### Genome coverage profiling

We used indexcov^10^ to estimate coverage from the Illumina whole genome sequencing library cross-site comparison data set. The bam file for each library used as input to indexcov^10^ to generate a linear index for each chromosome indicating the file (and virtual) offset for every 16,384 bases in that chromosome. This gives the scaled value for each 16,384-base chunk (16KB resolution) and provides a high-quality coverage estimate per genome. The output is scaled to around 1. A long stretch with values of 1.5 would be a heterozygous duplication; a long stretch with values of 0.5 would be a heterozygous deletion.

### Preprocessing and Alignment of 10X Genomics WGS Data

The 10X Genomics Chromium fastq files were mapped and reads were phased using LongRanger to the hg38/GRCh38 reference genome using the LongRanger v2.2.2 pipeline [https://genome.cshlp.org/content/29/4/635.full]. The linked-reads were aligned using the Lariat aligner^11^, which uses BWA MEM^3^ [Li H. et al. 2010] to generate alignment candidates, and duplicate reads are marked after alignment. Linked-Read data quality was assessed using the 10X Genome browser Loupe. MultiQC(v1.9)^7^ was run to generate an aggregated report in html format. A standard QC metrics report was generated from a custom script. The preprocessing and alignment QC analysis pipeline is described in **Suppl. Figure 1a**.

### Preprocessing and Alignment of Ion Torrent Data

Raw reads were first filtered for low-quality reads and trimmed to remove adapter sequences and low-quality bases. This step was performed using the BaseCaller module of the Torrent SuitTM software package v5.8.0 (Thermo Fischer Scientific Inc). Low-quality reads were retained from further analysis in the raw signal processing stage. Low-quality bases were trimmed from the 5’ end if the average quality score of the 16-base window fell below 16 (Phred scale), cleaving 8 bases at once. Processed reads were mapped to the GRCh38 reference genome by TMAP module of the Torrent Suite software package using the default map4 algorithm with recommended settings. Picard (v1.84)^5^ was then used to mark PCR and optical duplicates on the BAM files.

### Preprocessing and alignment for AmpliSeq

Low-quality bases and adapter sequences were trimmed with Trimmomatic^2^. The trimmed reads were mapped to the human reference genome GRCh38 (see the read alignment section) using BWA MEM (v0.7.17)^3^ in paired-end mode. We calculated on-target rate based on the percentage of mapped reads that were overlap the target capture bait region file (target.bed). We counted the number of variant-supporting reads and total reads for each variant position with MQ ≥ 40 and BQ ≥ 30 cutoffs. The preprocessing and alignment QC analysis pipeline is described in **Suppl. Figure 1a**.

### Somatic Variant Analysis

Four somatic variant callers, MuTect2 (GATK 3.8-0)^12^, SomaticSniper (1.0.5.0)^13^, Lancet (1.0.7), and Strelka2 (2.8.4)^14^, which are readily available on the NIH Biowulf cluster, were run using the default parameters or parameters recommended by the user’s manual. Specifically, for MuTect2, we included flags for “-nct 1 -rf DuplicateRead -rf FailsVendorQualityCheck -rf NotPrimaryAlignment -rf BadMate -rf MappingQualityUnavailable -rf UnmappedRead -rf BadCigar”, to avoid the running exception for “Somehow the requested coordinate is not covered by the read”. For MuTect2, we used COSMIC v82 as required inputs. For SomaticSniper, we added a flag for “-Q 40 -G -L –F”, as suggested by its original author, to ensure quality scores and reduce likely false positives. For TNscope (201711.03), we used the version implemented in Seven Bridges’s CGC with the following command, “sentieon driver -i $tumor_bam -i $normal_bam -r $ref --algo TNscope --tumor_sample $tumor_sample_name --normal_sample $normal_sample_name -d $dbsnp $output_vcf”. For Lancet, we ran with 24 threads on the following parameters “--num-threads 24 --cov-thr 10 --cov-ratio 0.005 --max-indel-len 50 -e 0.005”. Strelka2 was run with 24 threads with the default configuration. The rest of the software analyzed was run as a single thread on each computer node. All mutation calling on WES data was performed with the specified genome region in a BED file for exome-capture target sequences.

The high confidence outputs or SNVs flagged as “PASS” in the resulting VCF files were applied to our comparison analysis. Results from each caller used for comparison were all mutation candidates that users would otherwise consider as “real” mutations detected by this caller.

### GATK indel realignment and quality score recalibration

The GATK (3.8-0)-IndelRealigner was used to perform indel adjustment with reference indels defined in the 1000Genome project (https://www.google.com/url?sa=t&rct=j&q=&esrc=s&source=web&cd=4&ved=0ahUKEwjlkcfB5-nbAhVOhq0KHXUWCKUQFgg7MAM&url=ftp%3A%2F%2Fftp.1000genomes.ebi.ac.uk%2Fvol1%2Fftp%2Ftechnical%2Freference%2FGRCh38_reference_genome%2Fother_mapping_resources%2FALL.wgs.1000G_phase3.GRCh38.ncbi_remapper.20150424.shapeit2_indels.vcf.gz&usg=AOvVaw0pLCj6zDgJg0A6zbFeMfQl). The resulting BAM files were then recalibrated for quality with “BaseRecalibrator” and dbSNP build 146 as the SNP reference. Finally, “PrintReads” was used to generate recalibrated BAM files.

### Tumor Ploidy and clonality analysis from whole genome and exome data

To estimate the HCC1395 cell line ploidy, we used PURPLE^17^ to determine the purity and copy number profile. To determine the clonality of HCC1395 and HCC1395 BL, we performed somatic SNV and CAN analysis using superFreq^16^ on capture WES datasets. Mapped and markDuplicate bam files of a pair of HCC1395 and HCC1395BL were used as input and bam files of the remaining replicates of the HCC1395BL library were used to filter background. Analysis was run using the superFreq default parameters. The clonality of each somatic SNV was calculated based on the VAF, accounting for local copy number. The SNVs and CNAs undergo hierarchical clustering based on the clonality and uncertainty across replicates for the tumor sample.

### Assessment of reproducibility and O_Score calculation

We created and used “tornado” plots to visualize the consistency of mutation calls derived from aligners, callers, or repeated NGS runs. The height of the “tornado” represents the number of overlapping calls in the VCF files in descending order. The top of each plot portrays SNVs called in every VCF file. The bottom of the plots contains SNVs present in only one VCF file. The width of the “tornado” represents the number of accumulated SNVs in that overlapping category, which is scaled by the total number of SNVs in the corresponding sub-group. In addition, we established following formula to measure reproducibility based on the overlapping SNVs:

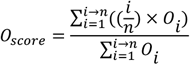

where n is the total number of VCF results in the pool set, i is the number of overlaps, *O*_*i*_ is the number of accumulated SNVs in the set with i number of overlapping.

## Data Records

All raw data (FASTQ files) are available on NCBI’s SRA database (SRP162370). The truth set for somatic mutations in HCC1395, VCF files derived from individual WES and WGS runs, and source codes are available on NCBI’s ftp site (ftp://ftp-trace.ncbi.nlm.nih.gov/seqc/ftp/release/Somatic_Mutation_WG/). Alignment files (BAM) are also available on Seven Bridges’ s Cancer Genomics Cloud (CGC) platform.

## Technical Validation

### Assessment of whole genome and exome sequencing data quality

For whole genome sequencing, fresh DNA samples were prepared using standard TruSeq PCR-free libraries prepared from 1000 ng input DNA. A total of 24 data sets were generated from six sequencing centers. There were three different Illumina sequencing platforms in the cross-platform comparison including HiSeq4000, HiSeq X Ten, and NovaSeq 6000. The quality assessment was based on the NGS preprocess pipeline produced quality metrics including Percentage of Q30 bases, sequencing yields, percentage of adapter sequences, percentage of mapped reads to reference genome, percentage of non-duplicate reads, GC content, DNA fragment insert sizes, genome coverage, etc (**Suppl. Figure 1**).

All sequencing centers and platforms produced high quality data as base call Phred quality scores above Q30, and greater than 99% of reads mapped to the reference genome (**Figure 2a**). The variation was observed in read coverage which was driven by sequencing platform yield differences as well as sequencing library pooling variations. Most sequencing sites produced genome coverage 50X (1,250 millions pair-end reads) per library, one sequencing site targeted about 100X (2,500 millions pair-end reads) per genome sequencing depth (**Figure 2b, Suppl. Figure 2a**). For whole exome sequencing, SureSelect Target Enrichment Reagent kit, PTN (Part No G9605A), SureSelect Human All Exon v6 and SureSelect Human All Exon v6 +UTRs were used, and sequencing was generated from 6 sequencing centers. Illumina Hiseq4000, Illumina Hiseq3000/4000, and Illumina Hiseq2500 were used. Sequencing quality from all sequences are high with greater than 99.1% of reads mapped to reference genome across sites. The variation was also observed in read coverage, most sequencing sites produced exome region on-target coverage 100X per library, and two sequencing sites targeted about 300X and 550X per genome sequencing depth (**Figure 2c**). When comparing WGS to WES libraries for the percentages of non-duplicated reads, all WGS libraries have consistently high percentages of non-duplicate reads, which indicates higher library complexity of WGS libraries than the targeted captures. In addition, there are much high variations in targeted exome capture libraries.

**Figure 2.**
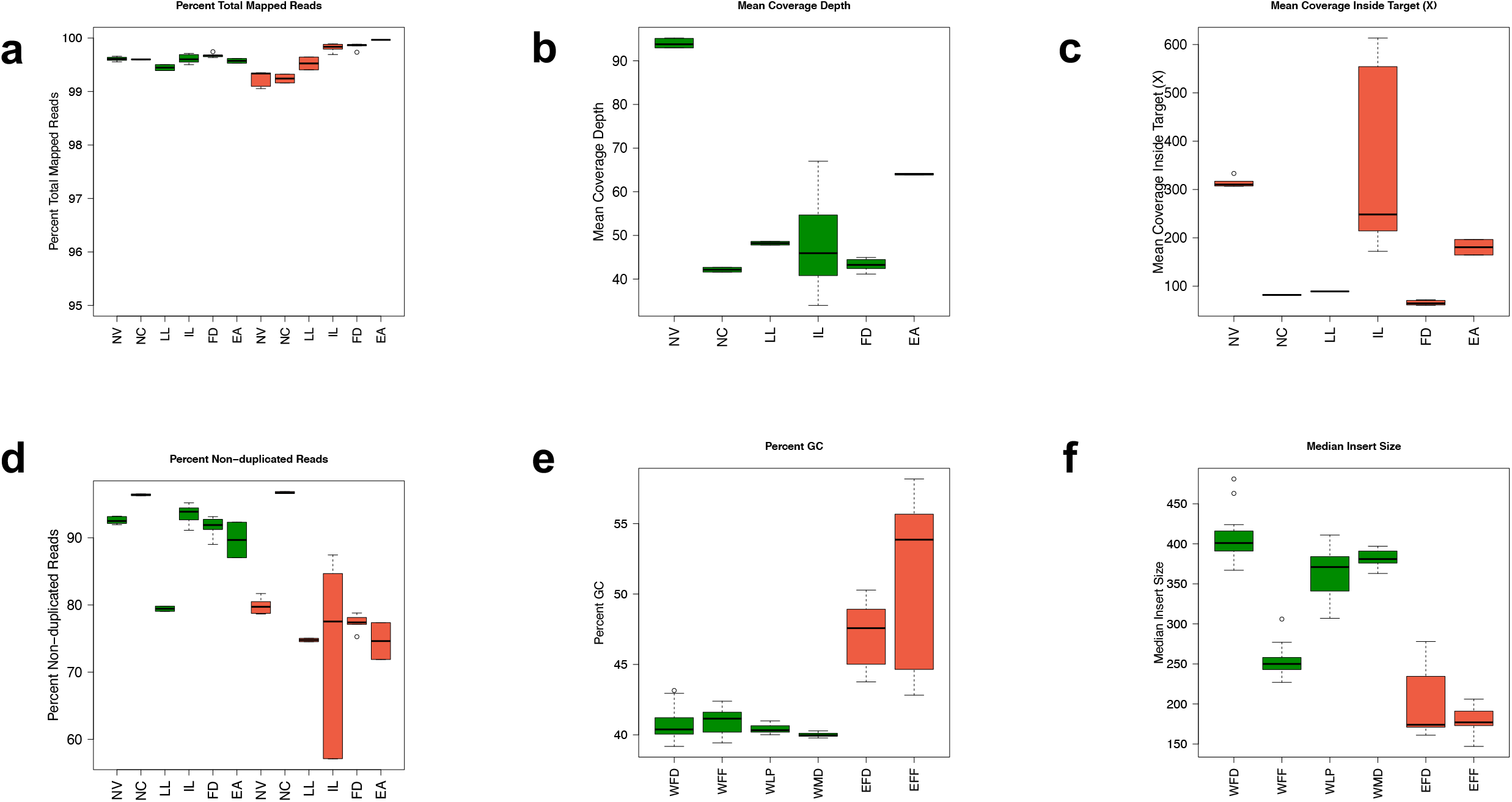
Overall data quality for WGS and WES data sets from Illumina platform. **(a)** Percentage of total reads mapped to reference genome (hg38) for WGS (Green) and WES (Red) across 6 sequencing sites. **(b)** Mean coverage depth for WGS libraries across 6 sequencing sites. (**c)** Mean coverage depth in target capture regions for WES libraries across 6 sequencing sites. **(d)** Percentage of non-duplicated reads mapped to reference genome across 6 sequencing sites. WGS (Green) and WES (Red). (**e)** Percent GC content from different library prep protocols. WGS (Green) and WES (Red). (**f)** Mean insert size distribution from different library prep protocols. WGS (Green) and WES (Red).

To determine if the quality of sequencing data was substantially different between different protocols, we also compared fresh DNA vs. FFPE DNA, different library protocols and input DNA amount, as well as mixture tumor DNA and normal DNA for profiling the tumor purity effect. Among the WGS libraries prepared using fresh cells, insert size distribution and G/C content were uniform (40 – 43% G/C). WES libraries have higher GC content (47.2% for fresh cells libraries, 51.1% for FFPE libraries) as well as higher variation (**Figure 2e**). All of the WGS libraries had very low adapter contamination (<0.5%) (**Suppl. Figure 2b)**, while WES libraries have higher adapter content due to smaller DNA fragment insert sizes (**Figure 2f**). WES library sizes are between 150bps -280bps for fresh cells. FFPE WGS libraries all have much shorter libraries sizes (225 - 300bps) than fresh DNA prepared WGS libraries (360 – 480bps). The libraries with higher adapter contamination also had much higher G/C content compared with the rest of the WES libraries (**Figure 2e**). When comparing library preparation kits across different DNA inputs across TruSeq PCR-free (1000ng), TruSeq-Nano, and Nextera Flex libraries prepared with 250, 100, 10, or 1 ng of DNA input, the percentage of non-redundant reads was very low (<20%) for TruSeq-Nano with 1 ng input, due to PCR amplification of a low input amount of DNA; higher input amount libraries have better performance; for the same input amount, Nextera Flex libraries have less variation and higher percentages of non-duplicated reads (**Suppl. Figure 2c**). We conclude the Nextera Flex library protocol might be a better option for low input DNA library preparation.

### Assessment of reference sample sequencing coverage and genome heterogeneity

We chose 26 replicates of HCC1395 and HCC1395BL data sets, which were libraries prepared using the Ilumina TruSeq DNA PCR free (1000ng) protocol and sequenced on Illumina HiSeq and NovaSeq. Each library was ranged from 50X to 100X genome coverage (**Figure 3a**). The percentage of genome coverage with less than 5X is 0.9 – 7.7% (**Suppl. Figure 4a**). For 10X Chromium libraries, each library has 45X - 120X genome coverage (**Figure 3b**), 6.4 – 7.3% of genome regions have read coverage less than 5X (**Suppl. Figure 4b**). 10X Chromium linked read technology produced input DNA molecule length in the range between 54 – 77kb. The site-to-site variation was due to sequencing depth differences. For WES samples, the target region has nearly 100% coverage by sequencing reads, however, we observed high variation in the sequencing coverage within each replicate as well as among replicates (**Suppl. Figure 3c**).

**Figure 3.**
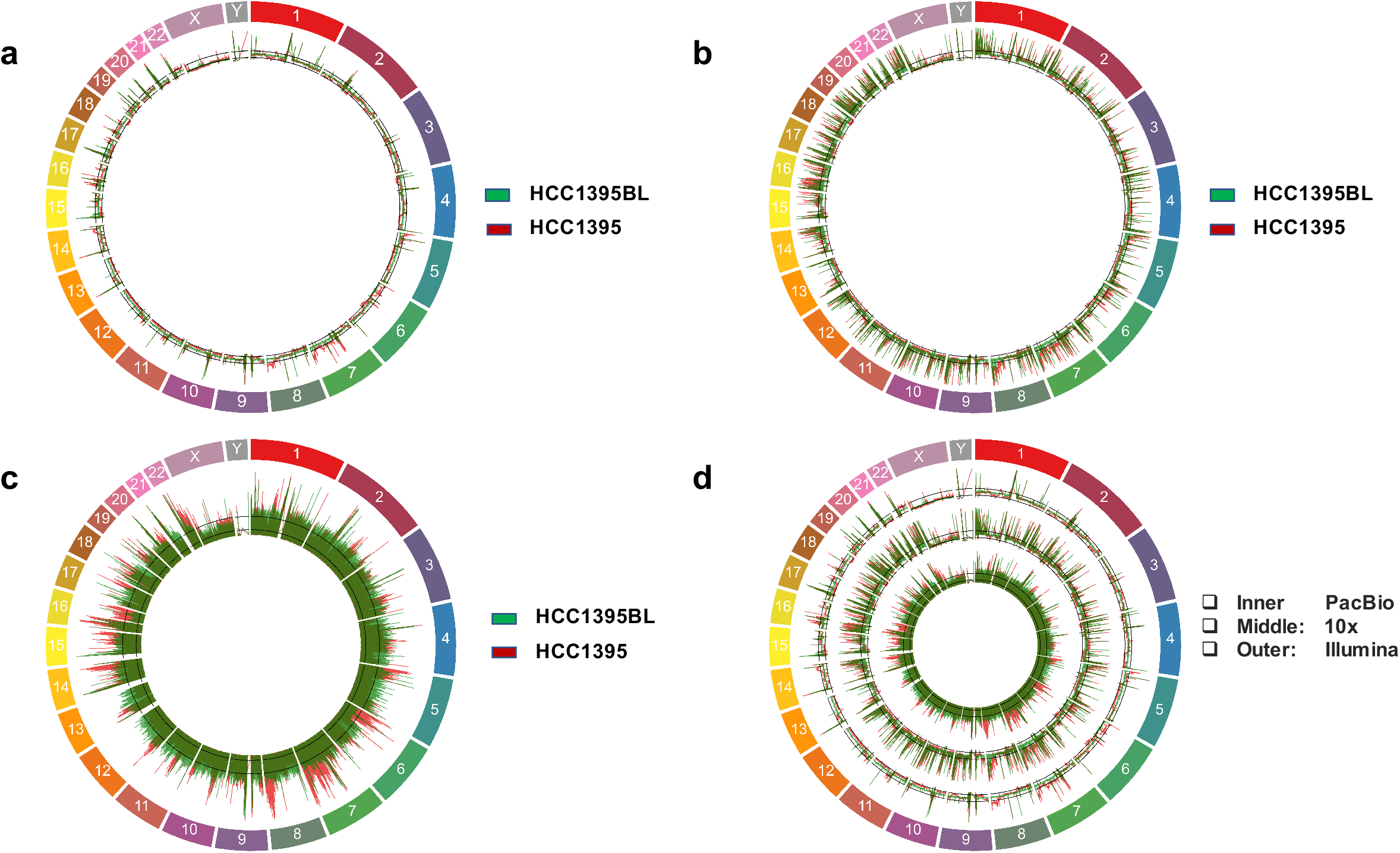
Genome coverage from WGS data from three technologies including Illumina, PacBio, and 10X Genomics. Outer rainbow color track: chromosomes, red track: HCC1395, green track: HCC1395BL. **(a)** Genome coverage from WGS data by reads from Illumina platform. **(b)** Genome coverage from WGS data by reads from 10X Chromium linked-read technology **(c)** Genome coverage from WGS data by reads from PacBio platform. **(d)** Genome coverage from WGS data by reads from 3 platforms together. Inner track: PacBio. Middle track: 10X Genomics. Outer track: Illumina.

**Figure 4.**
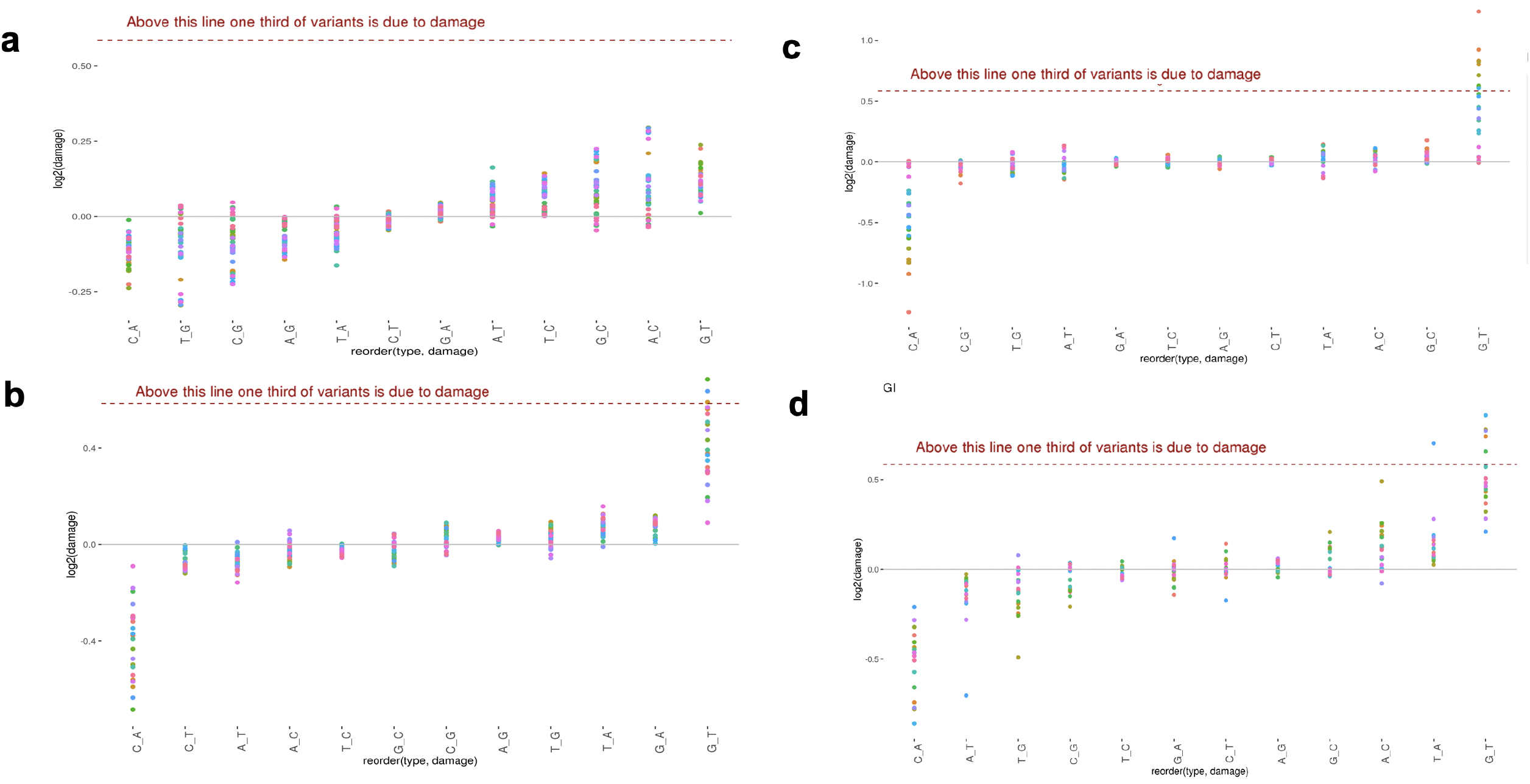
Evaluation of DNA damage for WGS and WES libraries. using GIV scores to capture the DNA damage due to the artifacts introduced during genomic library preparation. The estimation of damage is a global estimation based in an imbalance between R1 and R2 variant frequency. GIV score above 1.5 is defined as damaged. Undamaged DNA samples have a GIV score of 1. **(a)** DNA damage estimated for fresh cell prepared DNA for WGS Illumina libraries across different sites. **(b)** DNA damage estimated for FFPE WGS Illumina libraries. **(c)** DNA damage estimated for fresh cells prepared DNA for WES Illumina libraries across different sites **(d)** DNA damage estimated for FFPE WES Illumina libraries.

In addition, we generated two PacBio libraries with 40X of genome coverage from subreads. Long reads improve the map ability in repetitive genome regions where short-reads might fail to map correctly. PacBio long-read sequencing may cover the genomic regions where short reads cannot be mapped especially in the high GC/AT or low complexity genomic regions (**Figure 3c**). However, its higher sequencing error rate than short-read sequencing affects the accuracy for the low-frequency somatic mutation discovery. The variation in genome coverage might be due to differences in sequencing technologies (**Figure 3d**). From the study, short reads WGS has better uniform coverage compared to long reads. However, there is better coverage for certain genomic regions in long-read technologies; most noticeable are the highly repetitive regions, extreme GC regions, or around the centromere regions.

The Indexcov^10^ scaled read depth on reference genome for HCC1395 (**Suppl. Figure 4a**) and HCC1395BL (**Suppl. Figure 4b**) showed HCC1395 harboring many Copy Number Variation (gain or loss) events on every chromosome; HCC1395BL genome largely remains diploid except for chr6 and chr16 and chrX. It showed loss of a chrX and a net loss of one copy of the short-arm of chr6 and loss of one copy of the long-arm of chr16. Cytogenetic analysis with Affymetrix Cytoscan HD microarray confirms the Cytogenetic view of HCC1395 which harbors many copy numbers gains or losses; Cytogenetic view of HCC1395BL confirms the losses of chr6p, chr16q, and chrX (NBT-RA46164).

For HCC1395 cell line, the tumor purity and ploidy estimated from Illumina WGS data set (**Suppl. Figure 5a**) using PURPL^17^ software showed the tumor purity is 99% and the ploidy is around 2.85. Cell ploidy histogram from 10X Chromium single cell CNV data set (**Suppl. Figure 5b**) displayed the vast majority of cells form a peak around ploidy 2.8. The analysis of 1270 cells for HCC1395 from 10X Single Cell CNV data set also revealed numerous chromosome gains and losses events (**Suppl. Figure 5c**) consistently in sub-populations of cells, which confirmed HCC1395 is a heterogeneous cell line.

### Assessment DNA Damage Artifacts

A previous study has revealed that DNA damage accounts for the majority of the false calls for the so-called low-frequency (1-5%) genetic variants in large public databases^8^. The DNA damage directly confounds the determination of somatic variants in those data sets. The Global Imbalance Value (GIV) score is commonly used to measure DNA damage based on an imbalance between paired-end sequencing R1 and R2 variant frequency^8^. GIV scores to capture the DNA damage due to the artifacts introduced during genomic library preparation, the combination of heat, shearing, and contaminates can result in the 8-oxoguanine base pairing with either cytosine or adenine, ultimately leading to G>T transversion mutations during PCR amplification^18^. In addition, Formaldehyde also causes the deamination of guanine. FFPE is known to cause G>T/C>A artifacts^19^.

We calculated GIV score to monitor DNA damage in Illumina WGS and WES runs for both fresh DNA libraries as well as FFPE libraries. We found lower GIV scores for the G>T/C>A mutation pairs in fresh DNA WGS libraries (**Figure 4a)** than FFPE WGS libraries (**Figure 4b)**. In addition, both fresh cell DNA WES (**Figure 4c)** and FFPE WES Libraries (**Figure 4d)** all showed increased GIV scores for the G>T/C>A mutation pairs relative to WGS libraries. The GIV for G>T/C>A scores was inversely correlated with insert fragment sizes, and it is positively correlated to DNA shearing time (**Suppl. Figure 5a/b/c**); WES libraries have consistently shorter library insert sizes than all WGS library sizes (**Figure 2f)**. Thus, the GIV of G>T/C>A is a good indicator of DNA damage introduced during genomic library preparation. We observe the libraries have high G>T/C>A GIV scores also have a higher percentage of C/A mutation called in WES from private mutation calls which are not shared among replicates as displayed in **Suppl. Figure 5d**. Therefore, in order to improve cancer genomic variant call accuracy, effective mitigation strategies to improve library preparation methods, or software tools to detect and remove the DNA damage mutation calls are essential.

### Assessment reproducibility of somatic mutation calling from WES and WGS data sets

To assess the concordance and reproducibility of the somatic variant detection with both WES and WGS, we compared 12 replicates of WGS and WES for the matched tumor and normal cell lines carried out at six sequencing centers. Using three mutation callers (MuTect2^12^, Strelka2^13^, and SomaticSniper^14^) on alignments from three aligners (Bowtie2^15^, BWA MEM^3^, and NovoAlign), we generated a total of 108 variant call files separately. We were able to assess inter- and intra-centers reproducibility of the WES and WGS using the 12 repeat runs. The Venn diagram is widely used to display concordance of mutation calling results from a small number of repeated analyses; however, this type of diagram is not suitable for large data sets. To address this challenge, we applied the “tornado” plot to visualize the consistency of mutation calls. The number of SNVs unique to one single VCF file are represented by the width of the tornado at the bottom, and the number of SNVs called in all VCF files are represented by the width at the top. Thus, like the actual meteorological event, a tornado that is touching down is “bad” (many called variants are likely false positives), and a tornado with the majority of the data at the top is “better” (many common variants called across all conditions). As shown at the top of each plot (**Figure 5a**), we observed relatively more library-specific variants at the bottom of the WES tornado plots (bottom of tornado). In contrast, majority of called mutations (top of tornado) were shared across all 12 WGS (**Figure 5b**). Therefore, calling results from WES tended to have more inconsistent SNV calls (bottom of tornado) than those from WGS, indicating that WES results were less consistent than WGS results (**Figure 5a/b**). Here we also introduced the O_Score, a metric to measure reproducibility of repeated analyses (see Methods). O_Scores for WES runs were not only significantly lower than WGS runs, but also more variable (**Suppl. Figure 7a**). In addition, we measured reproducibility between replicates of WGS runs from both NovaSeq and HiSeq platforms to assess cross-platform variation. Both platforms were remarkably similar in terms of reproducibility, indicating that results from HiSeq and NovaSeq are comparable (**Suppl. Figure 7b**). Overall, we observed the cross-center and cross-platform variations for WGS were very small, indicating that all individual NGS runs, regardless of sequencing centers or NGS platforms, detected most “true” mutations consistently for WGS runs.

**Figure 5.**
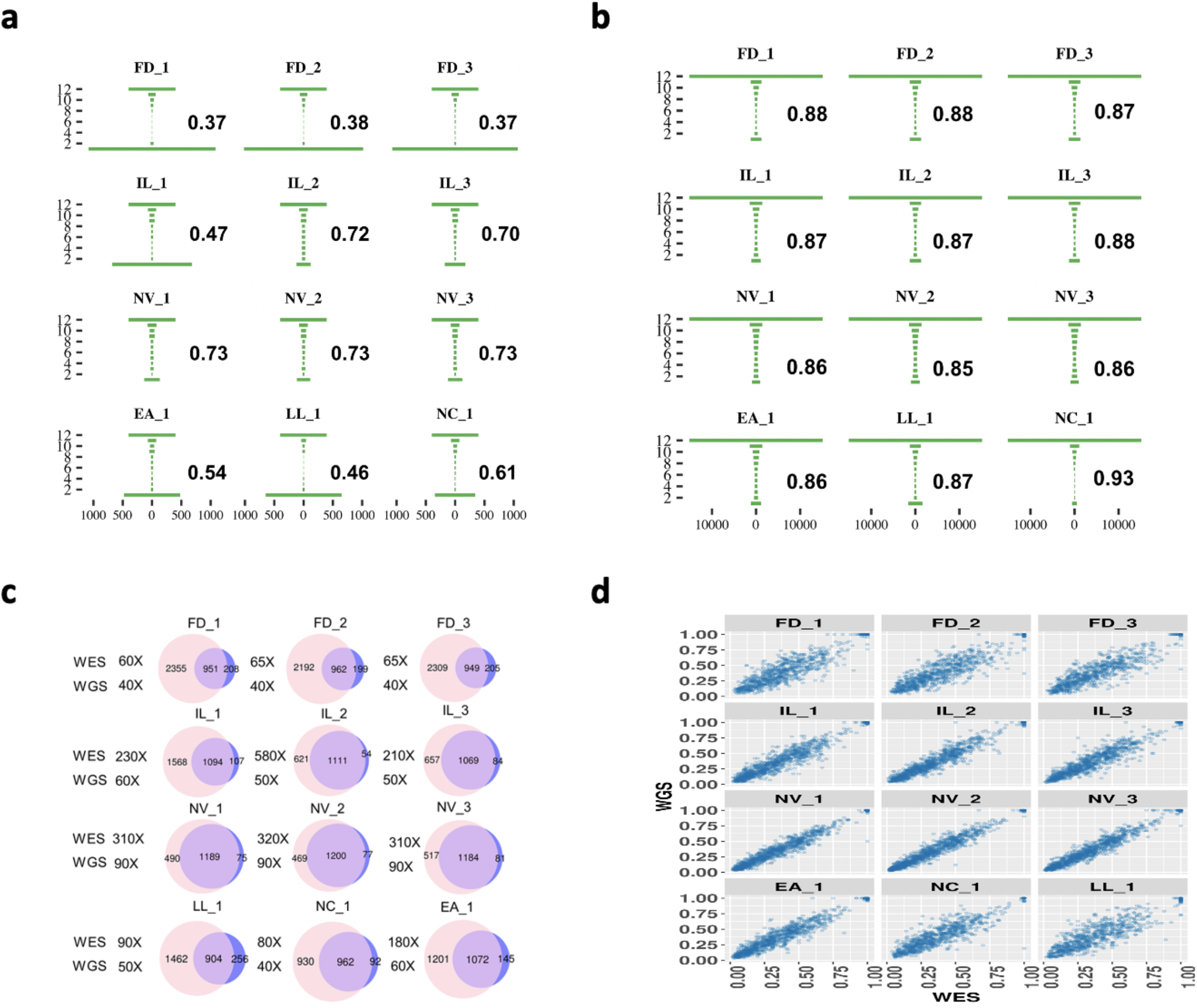
Reproducibility of somatic mutation calling from WES and WGS. The reproducibility “Tornado” plots for 12 repeated WES **(a)** and WGS runs **(b)**. The number in each plot represents the O_Score reproducibility measurement. **(c)** SNVs/indels calling concordance between WES and WGS from twelve repeated runs. For direct comparison, SNVs/indels from WGS runs were limited to genomic regions defined by an exome capturing kit (SureSelect V6+UTR). WES is shown on the left in the Venn diagram and WGS is on the right. Shown coverage depths for WES and WGS were effective mean sequence coverage on exome region, i.e. coverage by total number of mapped reads after trimming. **(d)** Correlation of MAF in overlapping WGS and WES SNVs/indels from repeated runs.

We also computed SNVs/indels calling concordance between WES and WGS from twelve repeated runs. For direct comparison, SNVs/indels from WGS runs were limited to genomic regions defined by an exome capturing protocol (SureSelect V6+UTR). WGS has a smaller number of private calls for each sample than WES **(Figure 5c)**. We observed the overlap between the WES and WGS improved as sequencing depth increased. Moreover, the correlation of MAF in overlapping WGS and WES SNVs/indels from repeated runs are positively correlated with higher sequencing depth **(Figure 5d)**. This indicates the benefit of high read coverage not only improves the detection sensitivity of mutations with low MAF, but also increases reproducibility of the calling sets. Overall, our results indicate the inter-center variations for WES were larger than inter-center variations for WGS, whereas the difference between intra-center variation between WES and WGS was not significant. As a result, WGS had much less inter-center variation and thus provided better reproducibility than WES for cancer genomic variants detection.

## Supporting information

Online-Only Supplemental Table 1-10

## Code Availability

All code used in processing the whole genome and exome-seq data are available on GitHub at the following link: https://github.com/abcsFrederick/NGS_Preprocessing_Pipeline

## Acknowledgements

The authors would like to thank Drs David Goldstein and Mariam Malik of the Office of Technology and Science at National Cancer Institute (NCI), National Institutes of Health (NIH), for the sponsorship and the usage of the NIH Biowulf cluster and support for this study; Dr. Jack Collins of the Advanced Biomedical and Computational Sciences, and Dr. Eric Stahlberg of Biomedical Informatics and Data Science Directorate at Frederick National Laboratory for Cancer Research for reviewing manuscript and providing suggestions; Seven Bridges Genomics for providing storage and computational support on the Cancer Genomic Cloud (CGC); this work also used the computational resources of Frederick Research Computing Environment (FRCE) at Frederick National Laboratory for Cancer Research. The authors sincerely thank members at NCI Sequencing Facility at Frederick National Laboratory for Cancer Research for their sequencing support for this study. This project has been funded in part with Federal funds from the National Cancer Institute, National Institutes of Health, under Contract No. HHSN261201800001I. The genomic work carried out at the Loma Linda University (LLU) Center for Genomics was funded in part by the NIH grant (S10OD019960), the Ardmore Institute of Health (AIH) grant (2150141), the American Heart Association (AHA) grant ((18IPA34170301), and Dr. Charles A. Sims’ gift to LLU Center for Genomics. Drs. L. Shi and Y. Zheng were supported by the National Natural Science Foundation of China (31720103909), the National Key R&D Project of China (2018YFE0201600), and Shanghai Municipal Science and Technology Major Project (2017SHZDZX01). The work carried out at Uppsala University was supported by grants from the Swedish Research Council (2017-00630, 2019-01976) and the Knut and Alice Wallenberg Foundation. Ene Reimann was supported by the European Union through the European Regional Development Fund (Project No. 2014-2020.4.01.15-0012).

## Disclaimer

This is a research study, not intended to guide clinical applications. The views presented in this article do not necessarily reflect current or future opinion or policy of the US Food and Drug Administration. The content of this publication does not necessarily reflect the views or policies of the Department of Health and Human Services. Any mention of commercial products is for clarification and not intended as endorsement.

## Author contributions

WX and YZ conceived and designed the study. YZ and WX drafted the manuscript. YZ, WX, LTF, CW, JN, UL and DM edited the manuscript. BT, JS, YK, CW, EJ, CL, KI, YTZ, LS, VP, MS, TH, EP, JD, PV, RM, DG, SK, ER, AS, JN, UL, ZC, and WC performed library construction and sequencing. YZ, XW, LTF, BZ, ZS, SC, KT and XFC performed bioinformatics data analyses. YZ and WX managed the project. All authors reviewed the manuscript. YZ finalized and submitted the manuscript.

## Competing interests

Li Tai Fang is employee of Roche Sequencing Solutions Inc. Erich Jaeger is employee of Illumina Inc. Virginie Petitjean and Marc Sultan are employees of Novartis Institutes for Biomedical Research. Tiffany Hung and Eric Peters are employees of Genentech (a member of the Roche group). All other authors claim no conflicts of interest.

## Notes

### Competing Interest Statement

The authors have declared no competing interest.

### Summary of Updates

Figure 2 revised, Author afflictions updated. Supplemental files updated

## References

1. Morash M, Mitchell H, Beltran H, Elemento O, Pathak J. The Role of Next-Generation Sequencing in Precision Medicine: A Review of Outcomes in Oncology. J Pers Med. 2018;8(3):30. Published 2018 Sep 17. doi:10.3390/jpm8030030

2. Bolger, A. M., Lohse, M. & Usadel, B. Trimmomatic: a flexible trimmer for Illumina sequence data. Bioinformatics 30, 2114–2120 (2014).

3. Li, H. Aligning sequence reads, clone sequences and assembly contigs with BWA-MEM. arXiv:1303.3997 [q-bio] (2013).

4. Babraham Bioinformatics - FastQC A Quality Control tool for High Throughput Sequence Data. Available at: https://www.bioinformatics.babraham.ac.uk/projects/fastqc/. (Accessed: 4th September 2018)

5. Picard Tools - By Broad Institute. Available at: http://broadinstitute.github.io/picard/. (Accessed: 23rd December 2017)

6. Okonechnikov, K., Conesa, A. & García-Alcalde, F. Qualimap 2: advanced multi-sample quality control for high-throughput sequencing data. Bioinformatics 32, 292–294 (2016).

7. Ewels, P. MultiQC: Aggregate results from bioinformatics analysis across many samples into a single report. Bioinformatics 32(19), 3047–8 (2016).

8. Chen, L., Liu, P., Evans, T. C. & Ettwiller, L. M. DNA damage is a pervasive cause of sequencing errors, directly confounding variant identification. Science 355, 752–756 (2017).

9. Heng Li, Minimap2: pairwise alignment for nucleotide sequences, Bioinformatics, Volume 34, 18, 15 September 2018, Pages 3094–3100, https://doi.org/10.1093/bioinformatics/bty191

10. Pedersen, B. et al. Indexcov: fast coverage quality control control for whole-genome sequencing, GigaScience, 6, 2017, 1–6 doi: 10.1093/gigascience/gix090

11. Bishara, A., Liu, Y., Weng, Z., Kashef-Haghighi, D., Newburger, D. E., West, R., Sidow, A., & Batzoglou, S. (2015). Read clouds uncover variation in complex regions of the human genome. Genome research, 25(10), 1570–1580. https://doi.org/10.1101/gr.191189.115

12. Cibulskis, K., Lawrence, M., Carter, S. et al. Sensitive detection of somatic point mutations in impure and heterogeneous cancer samples. Nat Biotechnol 31, 213–219 (2013). https://doi.org/10.1038/nbt.2514

13. Saunders, C. T. et al. Strelka: accurate somatic small-variant calling from sequenced tumor–normal sample pairs. Bioinformatics 28, 1811–1817 (2012).

14. Larson, D. E. et al. SomaticSniper: identification of somatic point mutations in whole genome sequencing data. Bioinformatics 28, 311–317 (2012).

15. Langmead, B., Trapnell, C., Pop, M. & Salzberg, S. L. Ultrafast and memory-efficient alignment of short DNA sequences to the human genome. Genome Biol. 10, R25 (2009).

16. Flensburg, C, Sargeant, T, Oshlack, A, Majewski, IJ (2020) SuperFreq: Integrated mutation detection and clonal tracking in cancer. PLOS Computational Biology 16(2): e1007603. https://doi.org/10.1371/journal.pcbi.1007603

17. Cameron, D., Baber, J., Shale, C., Papenfuss, A., Valle-Inclan, J., Besselink, M., Priestley, C. GRIDSS, PURPLE, LINX: Unscrambling the tumor genome via integrated analysis of structural variation and copy number, bioRxiv 781013; doi: https://doi.org/10.1101/781013

18. Costello, M. et al. Discovery and characterization of artifactual mutations in deep coverage targeted capture sequencing data due to oxidative DNA damage during sample preparation. Nucleic Acids Res 41, e67 (2013).

19. Do, H. & Dobrovic, A. Sequence Artifacts in DNA from Formalin-Fixed Tissues: Causes and Strategies for Minimization. Clinical Chemistry 61 (1), 64–71 (2015).

