## Supplemental Figures for "Whole Genome and Exome Sequencing Reference Datasets from A Multi-center and Cross-platform Benchmark Study"

The Somatic Mutation Working Group of the SEQC-II Consortium

### Supplementary Figures

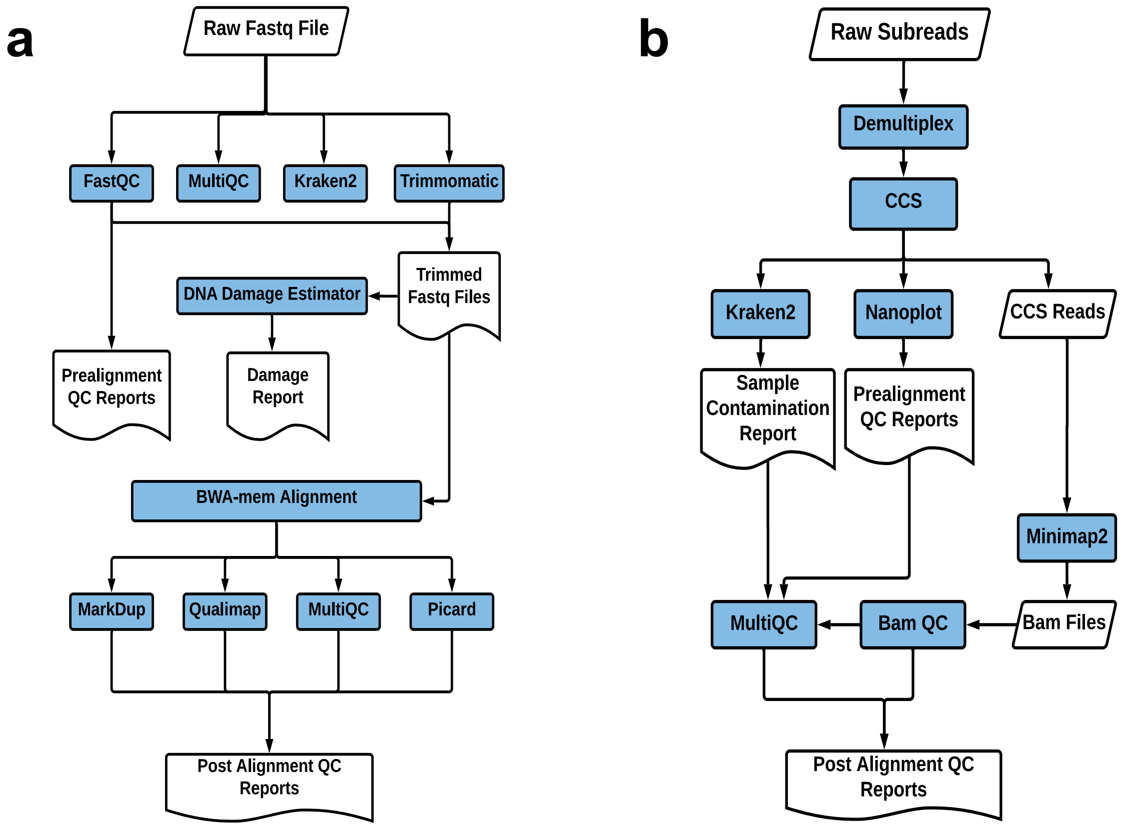


**Suppl. Figure 1:** Preprocessing and QC analysis pipelines used for whole genome and whole exome-seq data analysis **(a)** Short-read Illumina sequencing QC pipeline. **(b)** PacBio long-read sequencing QC pipeline.

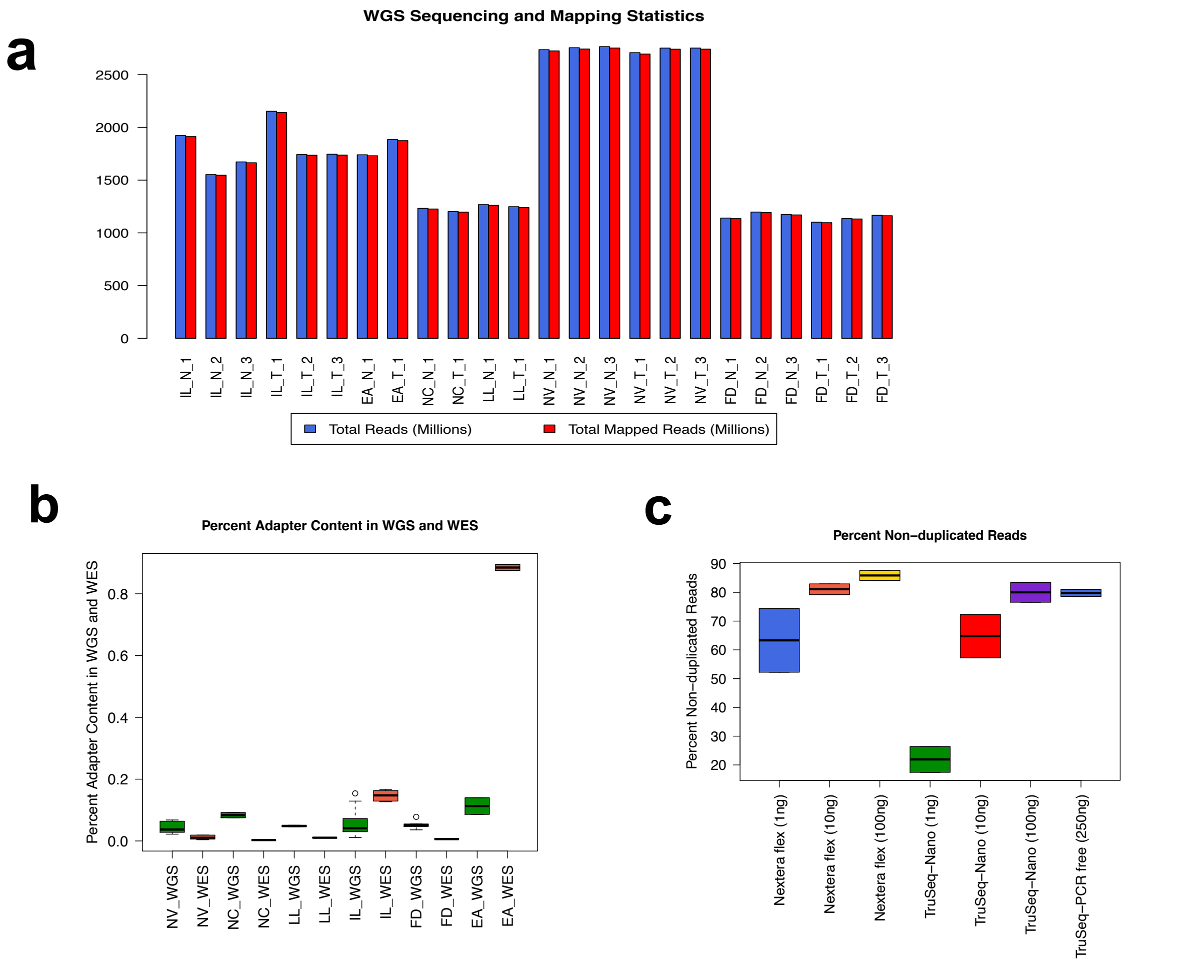


**Suppl. Figure 2:** WGS and WES cross-site data quality metrics. **(a)** WGS cross-site sequencing yields (Millions) mapped reads (Millions) statistics. **(b)** Adapter contents in WGS and WES Illumina short-read data set across 6 data centers **(c)** None duplicate mapped reads in Nextera Flex, TruSeq Nano and TruSeq PCR free protocols with different input amount.

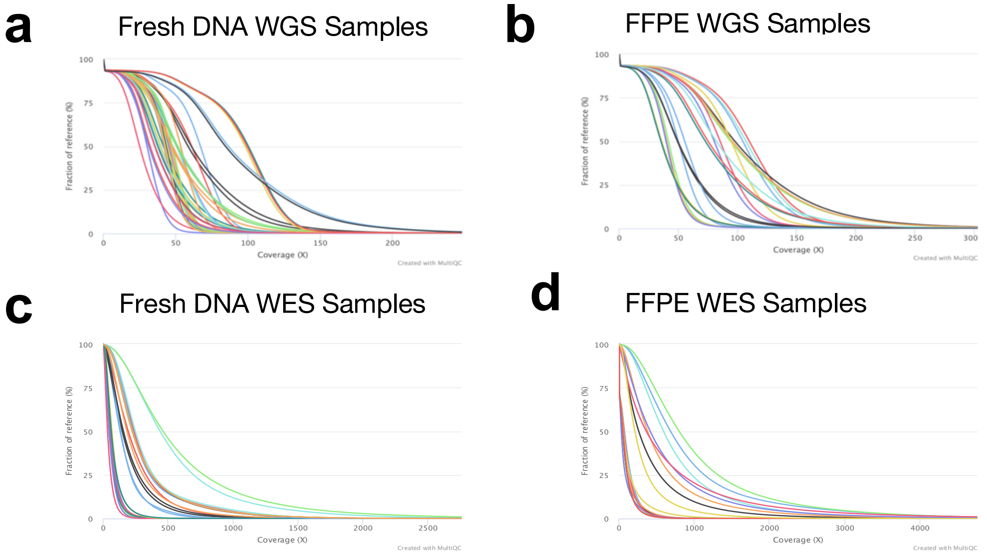

**Suppl. Figure 3:** Cumulative genome coverage for WGS and WES cross-site data sets and FFPE WGS and FFPE WES data sets. The graph displays the percentages of the reference genome with at least the given depth of coverage for each sample **(a)** Genome coverage for each fresh DNA prepared Illumina WGS libraries for cross-site comparison. **(b)** Genome coverage for FFPE WGS libraries. **(c)** Genome coverage for each Fresh DNA prepared Agilent SureSelect V6+UTR exome capture libraries for cross-site comparison. **(d)** Genome coverage for FFPE whole exome capture libraries.

**a**

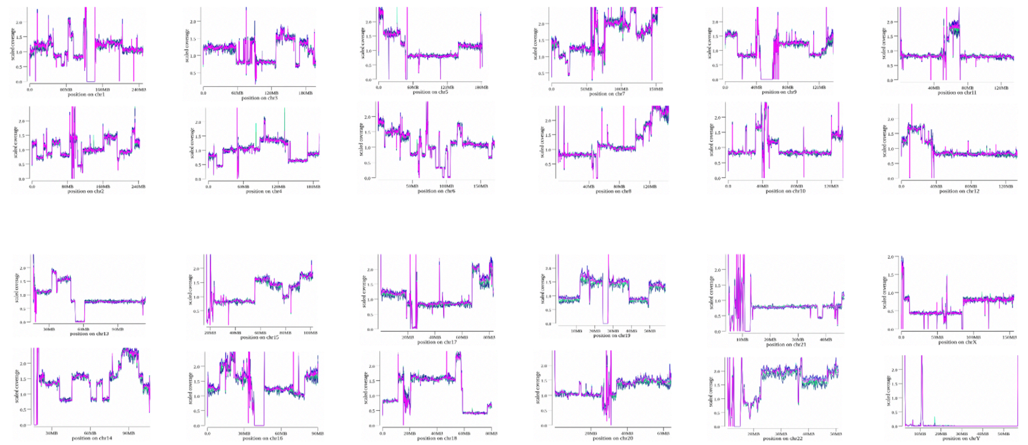

**b**

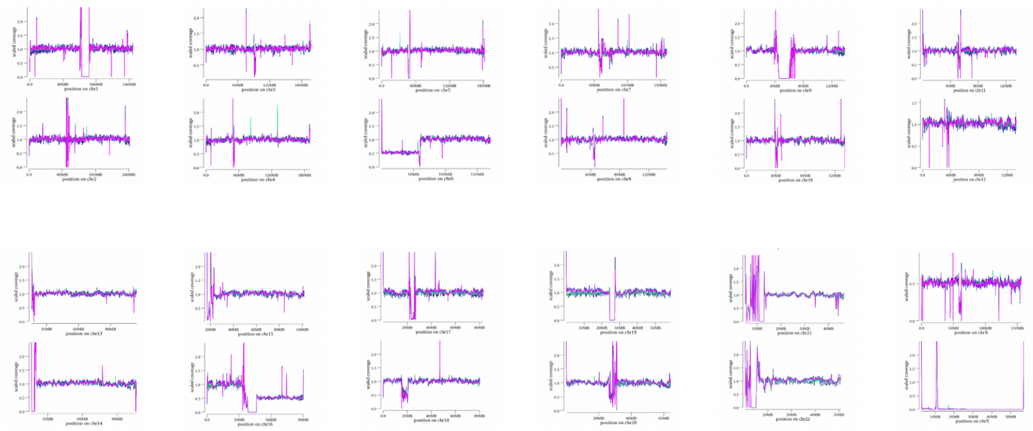

**Suppl. Figure 4:** Genome coverage plots generated using Indexcov software for whole genome sequencing cross-site comparison libraries. Indexcov gives the scaled value for each 16KB resolution and provides coverage values across the genome. A value of 1.5 would indicate a heterozygous duplication; a value of 0.5 would be a heterozygous deletion**. (a)** The estimated coverage along each chromosome for tumor HCC1395 cell line is shown. There are copy number gain or loss for each chromosome as shown in read coverage plot. **(b)** The estimated coverage along each chromosome for normal cell line HCC1395BL is shown. All chromosomes of HCC1395BL have the scaled index value of 1, which indicates the normal diploid genome except for chr6 and chr16 and chrX. It showed loss of a chrX and a net loss of one copy of the short-arm of chr6 and loss of one copy of the long-arm of chr16.

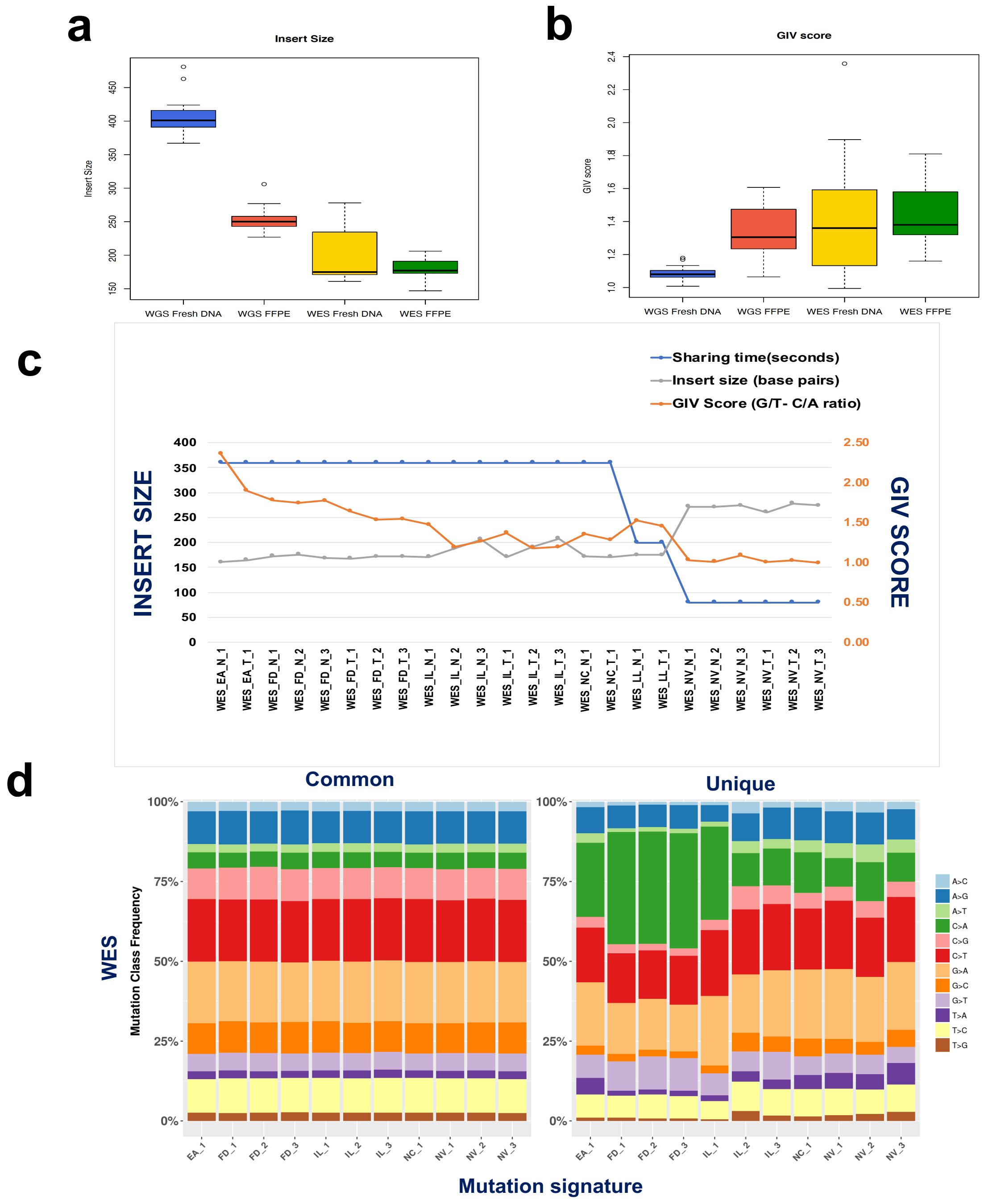


**Suppl. Figure 5:** Source of DNA damage artifacts and effect on cancer genome mutation calls. One type of DNA damage artifacts is introduced during DNA fragment mechanical shearing. Longer library inserts shave lower DNA damage scores (GIV scores) as shown in **(a)** library insert sizes for WGS cross-site libraries, WGS FFPE libraries, WES cross-site libraries, and WES FFPE libraries. **(b)** GIV score across different data sets; **(c)** correlation between library shearing time, insert sizes, and GIV scores. **(d)** percentage of mutation types for WES. The shared mutation across replicates is shown in left plot or sample specific unique mutations are shown in right plot. Note, high percentage of C/A mutation in the sites that also have high G/T_C/A GIV scores as displayed in **Suppl. Figure 5c.**

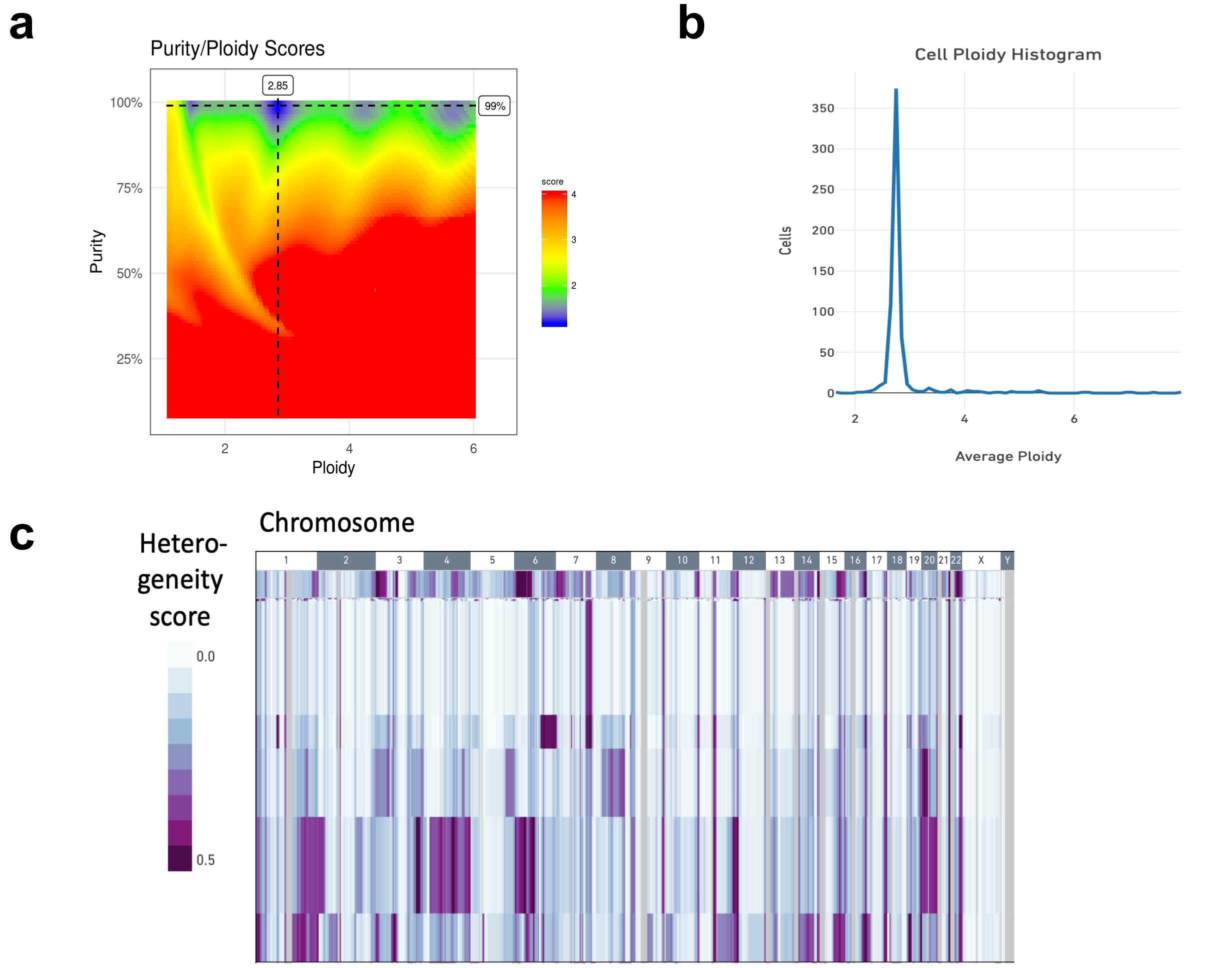


**Suppl. Figure 6:** HCC1395 tumor genome ploidy and heterogeneity measured from whole genome Illumina sequencing data and single cell CNV data. **(a)** Sample purity and ploidy estimated based on WGS for HCC1395 from Purple software, the tumor purity is above 99% with ploidy of 2.85. **(b)** Using 10X Genomics Single Cell CNV Solution, based on the analysis of 1270 cells for HCC1395 from 10x Single Cell CNV data set, Cellranger ploidy histogram displayed the vast majority of cells have ploidy of 2.8 as shown. **(c)** Heterogeneity analysis from10x Single Cell CNV data. Each row represents a cell being sequenced. Integer-scaled CNA profiles across the genome of 1270 HCC1395 cells were obtained. Similar cells were clustered together based on CNAs. Subclonal populations are marked in tracks. The chromosome-scale gains showed in darker purple, and losses displayed as light color in heatmap.

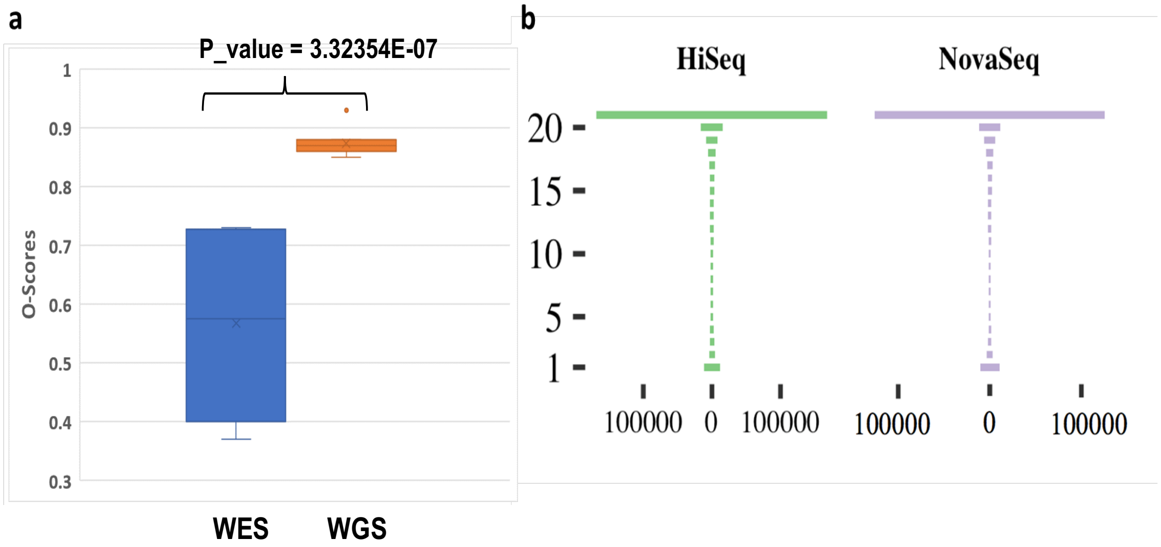


**Suppl. Figure 7:** Mutation calling repeatability and O_Score distribution. **(a)** Distribution of O_Score for 12 WGS and WES runs. **(b)** “Tornado” plot of reproducibility between 12 WGS runs on HiSeq and 9 WGS runs on NovaSeq (6000). SNVs/indels were called by Strelka2 on BWA alignments.
